# Dopaminergic signatures of state and trait-like pessimism in mice

**DOI:** 10.64898/2026.07.30.741818

**Authors:** Srijani Biswas, Steven J. Shabel

**Author notes:** Department of Neurobiology, University of California, San Diego, La Jolla, CA, 92093.

## Abstract

The midbrain dopaminergic system is critical for learning reward expectations, but how it contributes to pessimistic bias in reward expectation remains unclear. Using a Pavlovian ambiguous-cue task while recording mesolimbic dopaminergic activity, we find that trial-to-trial, “state” pessimism is associated with decreased dopaminergic activity during ambiguous cues, followed by a self-correcting positive shift in the balance of responses to reward and reward omission. Mice with pessimistically skewed cue interpretation, or “trait-like” pessimism, also had decreased activity during ambiguous cues. However, opposite of state pessimism, trait-like pessimism was associated with a negative shift in outcome signaling driven by larger reward-omission responses. These data indicate that state and trait-like pessimism arise from distinct neural mechanisms and support a model in which trait-like pessimism is maintained by asymmetrically weighted dopaminergic outcome signals.

## Introduction

Mesolimbic dopaminergic neurons are strongly implicated in learning reward expectations^1–13^. Phasic dopamine signals have been linked to reward prediction errors (RPEs) and temporal-difference (TD) reinforcement learning, providing a teaching signal that updates future expectations^1,2,4–9,14–18^. Within this framework, reward expectation is reflected in dopaminergic responses to learned cues, as well as, reward delivery and reward omission: increasing responses to learned cues and homeostatically decreasing responses to reward delivery and amplifying “dips” during reward omission that drive learning. Dopaminergic projections to the nucleus accumbens are particularly well positioned to influence motivation and reinforcement by integrating predictive and outcome information^15,19^.

In practice, reward expectations are not solely dictated by objective reward probabilities, especially under ambiguity, when cues only partially match learned predictors and animals must generalize, integrate prior beliefs, and weigh uncertain evidence^20,21^. Such belief-state computations are relevant to dopamine signaling because dopamine RPEs can reflect inferred state probabilities under uncertainty, not only the objective reward associated with an observable cue^22–24^. Under these conditions, the same cue can evoke different subjective expectations from one moment to the next and can be interpreted differently across individuals, such as in pessimism, a bias implicated in mood disorders^25,26^. Yet most paradigms quantify expectation coarsely or average across trials and subjects, limiting the ability to resolve how moment-to-moment subjective expectation maps onto dopaminergic signaling.

Despite extensive work on dopaminergic prediction errors and extensive work on pessimistic bias in affective science, the relationship between pessimistic expectation biases and dopaminergic dynamics remains unresolved. Although dopamine modulates affective bias^27^, it is unclear whether transient, trial-to-trial shifts within an individual (“state” pessimism) and stable differences between individuals (“trait-like” pessimism) are associated with similar changes in dopaminergic signals as decreasing the reward probability of a cue: reduced activity during the cue and a compensatory, positive shift in reward and reward omission signals. A positive shift in outcome signals during persistent, trait-like pessimism would be consistent with a model wherein trait-like pessimism is maintained by negatively shifted, asymmetric plasticity rules downstream of dopamine neuron activity^28^. However, symmetric or negatively shifted outcome responses during persistent, trait-like pessimism would favor a model wherein trait-like pessimism is maintained by asymmetric, negatively shifted weighting of RPE signals during reward and reward omission^29^.

Here, we address these questions by combining a Pavlovian ambiguous-cue task that yields a continuous, trial-resolved behavioral measure of reward expectation with recordings of dopaminergic projections to the nucleus accumbens. We operationalize state pessimism as trial-to-trial reductions in behavioral expectation within an animal during ambiguous cues, capturing transient subjective bias. We operationalize trait-like pessimism as a stable between-animal skew in ambiguous-cue interpretation across sessions, capturing persistent individual differences in bias.

Using this approach, we find that both state and trait-like pessimism are associated with reduced dopaminergic activity during ambiguous cues, consistent with a shared reduction in cue-evoked signaling during pessimistic interpretation. However, the two forms of pessimism diverge in outcome processing. Subjective, trial-to-trial pessimism is accompanied by a compensatory positive shift in the balance of outcome signals—enhanced dopamine responses during reward consumption and reduced RPEs during reward omission—consistent with adaptive TD updating that reduces the impact of negative outcomes while increasing sensitivity to positive outcomes when expectations are low. In contrast, mice with trait-like pessimistic interpretation exhibit a negative shift in the balance of outcome signals due to an increased omission-evoked RPE. Together, these findings support a model in which state pessimism engages flexible, homeostatic outcome updating, whereas trait-like pessimism is associated with a negatively shifted balance of positive and negative outcome signals that would sustain pessimistic expectations.

## Results

### Mesolimbic dopamine tracks learned reward expectations in a Pavlovian ambiguous-cue task

To determine how pessimism affects mesolimbic dopaminergic responses to reward-predictive cues, we expressed calcium indicators (jGCaMP8f or RCaMP1b) in VTA dopaminergic neurons and implanted optical fibers in the nucleus accumbens core to record mesolimbic dopaminergic activity (DA→NAc) during a Pavlovian ambiguous-cue task (Fig. 1A-C; Supplementary Figure 1A-D). Mice learned that an odor was followed by sucrose reward half the time, whereas odor– tone compound cues predicted either certain reward (positive tone; 3 or 12 kHz) or certain reward omission (negative tone; 12 or 3 kHz; Fig. 1B). The odor was critical for measuring bidirectional changes in licking related to reward expectation and for controlling for motivation and attention. During testing, intermediate-frequency ambiguous tones were presented and reinforced in proportion to their acoustic similarity to the positive/negative tones to prevent distortion of cue significance over time (Fig. 1B,C). All mice learned to lick during the odor and most mice showed discriminatory anticipatory licking during the positive and negative tones (hereafter, these tone learner mice are called “learner” mice, 19 of 28 mice, Fig. 1D and Supplementary Figure 2A,C; “non-learner” mice, 9 of 28; Supplementary Figure 2B,D; see Supplementary Table 1 for statistics). Learners had increased DA→NAc during the positive tone and decreased DA→NAc during the negative tone (Fig. 1E and Supplementary Figure 2E,G,I; Supplementary Table 1), consistent with previous experiments^30^. Non-learners had no difference in DA→NAc during the tone cues (Supplementary Figure 2F,H,J; Supplementary Table 1), consistent with changes in DA→NAc encoding the valenced motivational significance of the cues^31–33^. Learners also showed intermediate anticipatory licking and DA→NAc during the ambiguous cues (Fig. 1D,E; Supplementary Table 1), as expected.

**Figure 1.**
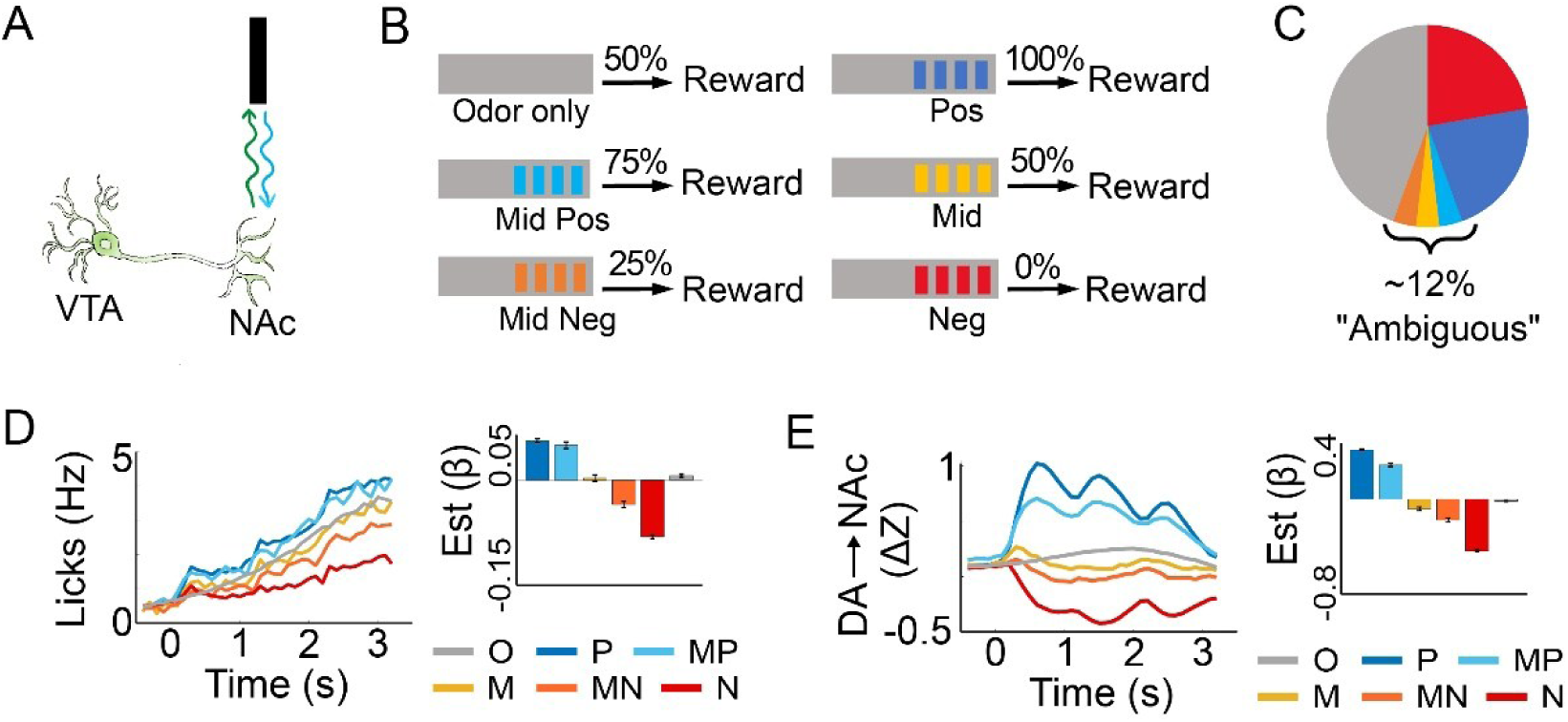
Licking and mesolimbic dopaminergic activity in the Pavlovian ambiguous-cue task. **(A)** Schematic showing fiber photometry of mesolimbic dopaminergic activity (DA→NAc) in DAT-cre mice. VTA, ventral tegmental area. NAc, nucleus accumbens. **(B)** Schematic of the cues and reward probabilities in the test sessions. Gray bars indicate 6 s of odor presentation. Colored bars indicate 200 ms pips of each tone. **(C)** Pie chart showing the proportion of each trial type in the test sessions. Colors as in B. **(D)** Left, mean licking in learners. Right, corresponding beta estimates and standard error from linear mixed-effects modeling (LME, see Methods). Zero on the x-axis indicates start of tones. O, odor only. P, positive tone. MP, mid-positive tone. M, mid tone. MN, mid-negative tone. N, negative tone. **(E)** Same as D, but showing DA→NAc instead of licking.

### DA→NAc encodes subjective state pessimism during ambiguous cues

Although on average learners showed intermediate anticipatory licking during the ambiguous cues, there was substantial variation in anticipatory licking to all cues (Supplementary Figure 3A,B). We hypothesized that some of the variation in anticipatory licking was due to transient changes in subjective expectation of reward (“state optimism”) and nonreward (“state pessimism”) and would be encoded in DA→NAc. To test this possibility, we split individual trials into optimistic and pessimistic categories, depending on the amount of anticipatory licking in each trial. To control for potential changes in attention and motivation over time, optimistic and pessimistic trials were defined as tone trials with more, or less, anticipatory licking than recent odor-only trials (Fig. 2A,B; Supplementary Figure 3C,E,F; Supplementary Table 1). Consistent with the hypothesis, optimistic trials had larger DA→NAc activity during ambiguous cues than pessimistic trials in learners (Fig. 2C; One-way RM ANOVA, *P* = 4.8 × 10^−5^; Supplementary Figure 3G; Supplementary Table 1). To maximize statistical power while controlling for possible correlations in data from individual mice and recording sites, we used linear mixed effect (LME) models that included data from each trial and modeled individual mice and recording sites as random effects^34^. This statistical strategy also showed that DA→NAc was larger during cues on optimistic than pessimistic trials, even when controlling for tone types which have different objective reward probabilities (LME, *P* = 1.8 × 10^−15^; Supplementary Table 1). The difference in DA→NAc during cues on optimistic and pessimistic trials is difficult to explain by lack of attention or motivation on pessimistic trials because activity decreased during pessimistic trials relative to odor-only trials (Fig. 2C, *P* = 5.0 × 10^−15^; Supplementary Table 1); if mice were not pessimistic but simply not attending as much or as motivated on pessimistic trials, DA→NAc should have remained constant or increased slightly during these trials. We also found that non-learners did not show the same difference in DA→NAc on optimistic and pessimistic trials (Supplementary Figure 3D,H; *P* = 0.82; Supplementary Table 1) despite similar differences in licking on optimistic and pessimistic trials as learners (Supplementary Figure 3C,F; Supplementary Table 1), indicating that differences in DA→NAc during ambiguous cues on optimistic and pessimistic trials in learners are also not due to differences in motor activity or trial engagement. The specificity of differences in DA→NAc on optimistic and pessimistic trials for learners also indicates that differences in DA→NAc during cues on optimistic and pessimistic trials are due to optimistic and pessimistic interpretation of environmental cues, not just optimism or pessimism per se.

**Figure 2.**
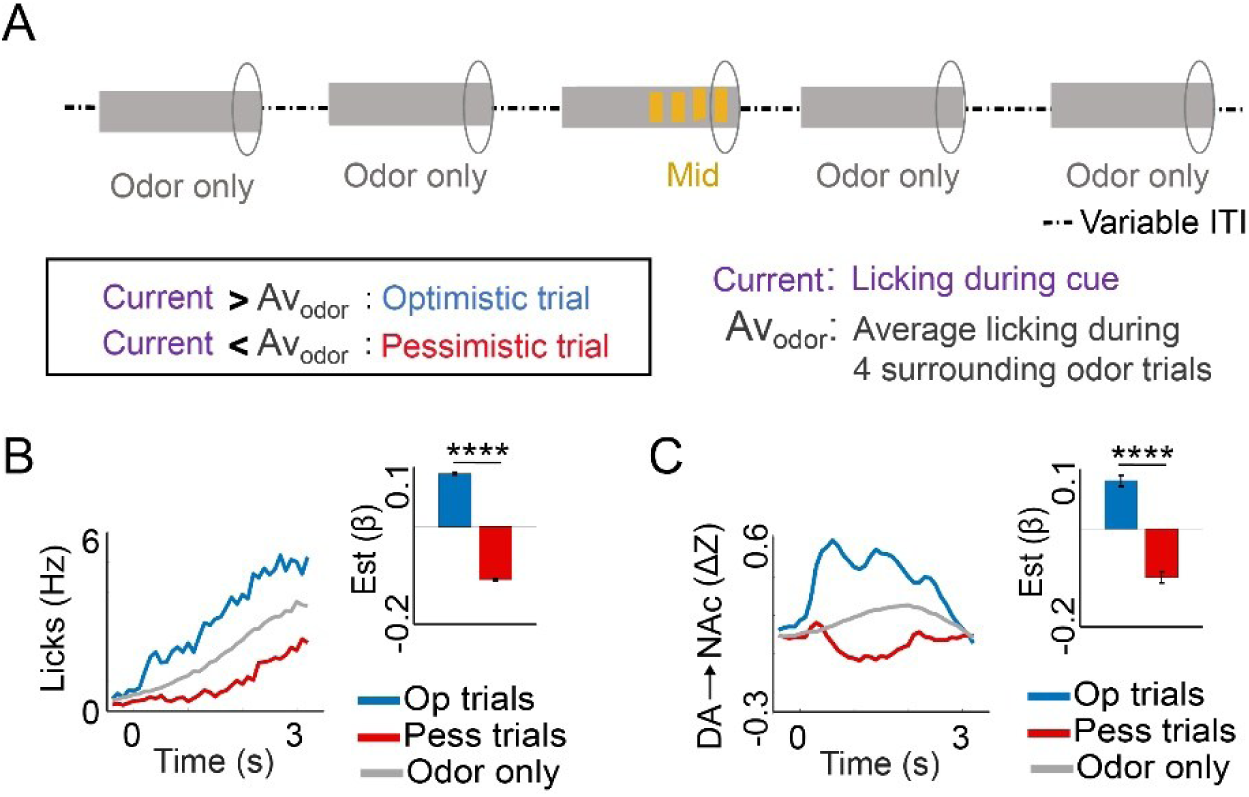
DA→NAc encodes state pessimism during ambiguous cues. **(A)** Schematic detailing the method used to categorize trials as pessimistic or optimistic, with surrounding odor trials used as reference. Circles indicate the time used for the trial classification. **(B)** Left, mean licking during odor-only trials (Odor) and optimistic (Op) and pessimistic (Pess) ambiguous cue trials. Right, corresponding beta estimates from LME which controls for objective trial type and possible correlations within mice and/or recording sites. Zero on the x-axis indicates start of tones. Data from 18 learner mice. **(C)** Same as B, but showing DA→NAc instead of licking. **** P < .0001. Please see Supplementary Table 1 for more statistical information.

We hypothesized that pessimistic trials might be more common immediately after non-rewarded trials. However, this was not the case (Supplementary Figure 4A, *P* = .42; Supplementary Table 1). There was also no significant effect of the previous trial when including all trial types (Supplementary Figure 4B, *P* = .062; Supplementary Table 1). Nevertheless, it remained possible that previous trial history, extending beyond the immediately preceding trial, would determine the likelihood of a pessimistic trial. To test this possibility, we took advantage of the fact that mice were often tested simultaneously in pairs with identical trial-type sequences. If pessimistic trial likelihood were greatly affected by trial-type history, then paired mice should have similarly low levels of anticipatory licking as reference mice during the reference mouse’s pessimistic trials (and high levels of anticipatory licking during the reference mouse’s optimistic trials; Supplementary Figure 4C). However, licking was very different between reference and paired mice on these trials (Licking: Supplementary Figure 4D,E; ref_paired_mouse x op_pess_trial interaction, *P* = 1.5 × 10^−68^; op trials, ref vs paired mouse, *P* = 2.8 × 10^−42^; pess trials, ref vs paired mouse, 7.1 × 10^−30^; Supplementary Table 1) and the difference in licking on optimistic and pessimistic trials in the paired mice did not reach statistical significance (paired mice, *P* = .055; reference mice, *P* << .0001, Supplementary Table 1). Similarly, DA→NAc on optimistic and pessimistic trials in the paired mice was not significantly different (Supplementary Figure 4F; paired mice, *P* = .76; reference mice, *P* = 9.4 × 10^−4^; Supplementary Table 1), indicating that optimistic and pessimistic trial classifications were largely subjective.

### State pessimism is associated with a compensatory positive shift in dopaminergic outcome signals

Because reward expectation is thought to decrease DA→NAc during the outcome period (causing RPEs) ^1,2,4–7,9,14,15^, our hypothesis further predicted a difference in the balance of positive and negative outcome encoding in DA→NAc during optimistic and pessimistic trials, with pessimism homeostatically tipping the balance toward positive outcome encoding in DA→NAc. Our data were consistent with this prediction (Fig. 3A, *P* = .0002; Supplementary Table 1). The altered balance during state pessimism was due to both decreased DA→NAc reward-omission responses (smaller magnitude changes) and increased DA→NAc reward responses (Sucrose omission: DA→NAc, Fig. 3B, *P* = 6.7 × 10^−15^; Sucrose consumption: DA→NAc, Fig. 3C, *P* = 1.9 × 10^−14^; objective reward probabilities associated with different tone types are accounted for by the LME; Supplementary Table 1), as expected. Together, these data indicate that DA→NAc encodes trial-to-trial subjective optimistic and pessimistic interpretation of cues (state optimism/pessimism), not simply objective reward probabilities.

**Figure 3.**
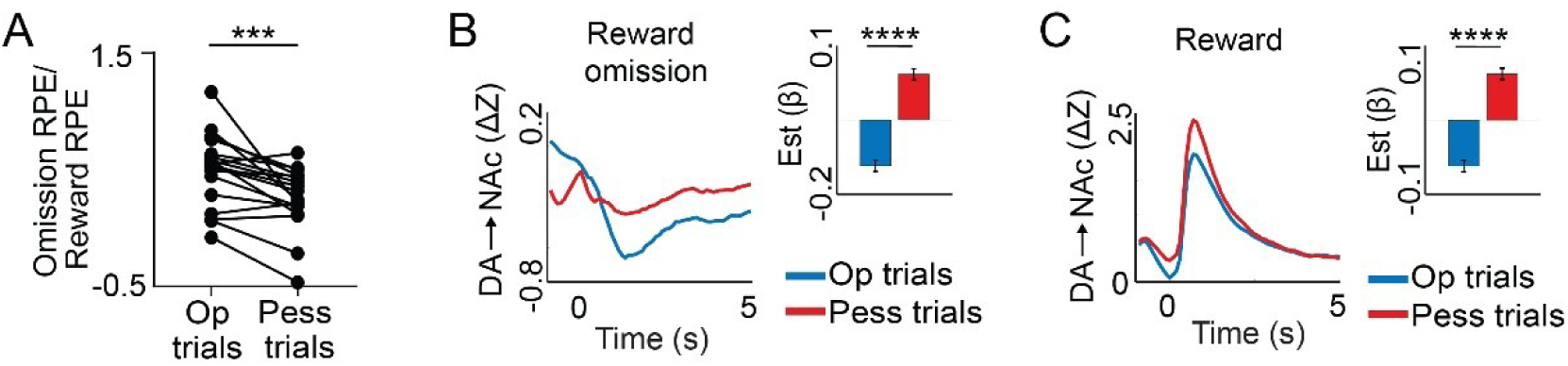
Positively shifted balance of dopaminergic outcome signals during state pessimism. **(A)** DA→NAc during sucrose omission relative to DA→NAc during sucrose consumption. Each pair of data points is from one learner mouse (N = 18). The sign of the omission activity was inverted to make the ratios more intuitive. **(B)** Left, mean DA→NAc during sucrose omission on optimistic (Op) and pessimistic (Pess) tone trials. Right, corresponding beta estimates from LME which controls for objective trial type and possible correlations within mice and/or recording sites. Zero on the x-axis indicates start of sucrose omission. Data from 18 learner mice. **(C)** Same as B, but during sucrose consumption. *** P < .001. **** P < .0001. Please see Supplementary Table 1 for more statistical information.

### Ambiguous-cue response skew reveals stable trait-like optimism and pessimism

Given the subjectivity in optimistic and pessimistic trial classifications, we hypothesized that there would also be consistent optimistic and pessimistic subjective bias among individual mice (trait-like optimism/pessimism). We used response skew during the ambiguous cues, as others did previously in ambiguous-cue paradigms^35^, as a measure of individual affective bias in learner mice (Fig. 4A-C). This measure accounts for individual differences in anticipatory licking rates, and is therefore theoretically independent from motivational state. Consistent with this, although anticipatory licking rates declined within sessions (Fig. 4D; *P* = .0002; Supplementary Table 1), anticipatory licking skew did not change systematically within sessions (Fig. 4E; *P* = .75; Supplementary Table 1). Additionally, there was no relationship between individual differences in overall anticipatory licking rates (average of all trial types) and individual differences in licking skew (Fig. 4F; *P* = .79; Supplementary Table 1). Importantly, licking skew of individual mice was stable across testing days, consistent with a trait-like measure (Fig. 4G, *r* = .83, *P* = .00002; Supplementary Table 1).

**Figure 4.**
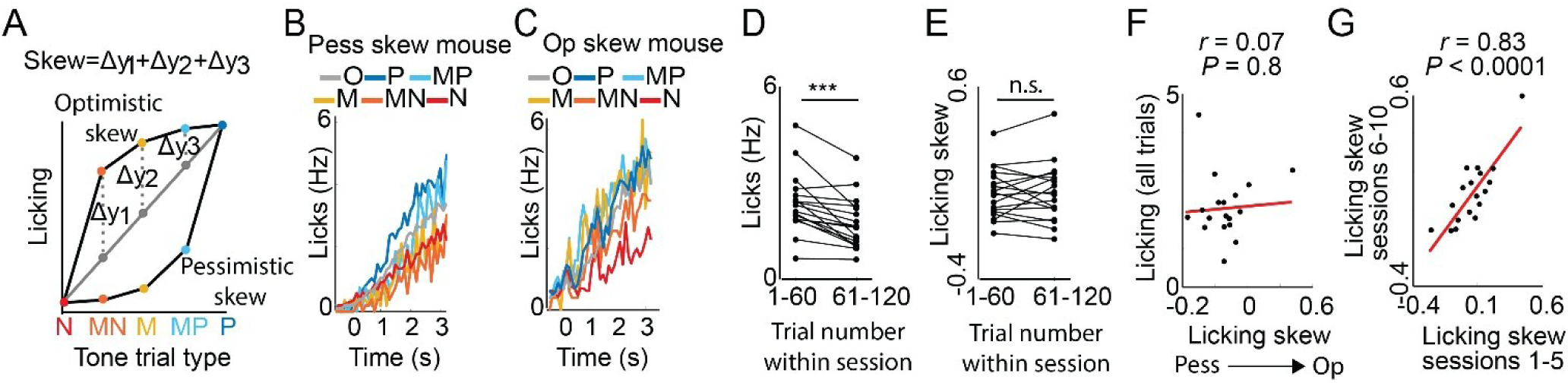
Licking skew during ambiguous cues as a measure of trait-like optimism/pessimism. **(A)** Schematic showing the calculation of licking skew. Expected licking values (gray dots) are calculated using linear interpolation (solid gray line) of mean licking responses for negative (N) and positive (P) tones. The differences between actual and expected mean licking for each ambiguous tone (dashed gray lines) are then added to calculate skew. Negative skews are “pessimistic” and positive skews are “optimistic”. **(B)** Mean licking during test sessions of a mouse with a pessimistic licking skew. Trial-type abbreviations as in Figure 1. Zero on the x-axis is the start of the tones. **(C)** Same as B, but in a mouse with an optimistic licking skew. **(D)** Mean anticipatory licking for all trials in the early (1-60) vs late (61-120) parts of the test sessions (each pair of data is from one mouse; N = 18 learner mice). **(E)** Same as D but showing licking skews instead of licking rates. **(F)** Mean anticipatory licking for all trials vs licking skew (N = 18 learner mice). **(G)** Licking skew in the last 5 vs first 5 test sessions (N = 18 learner mice). *** P < .001. Please see Supplementary Table 1 for more statistical information.

### Trait-like pessimism is encoded by reduced DA→NAc responses to ambiguous cues

To determine if DA→NAc during ambiguous cues encodes trait-like pessimism, we calculated DA→NAc skew, like licking skew (examples of DA→NAc in mice with pessimistic and optimistic licking skews in Fig. 5A,B). Consistent with trait-like optimism/pessimism encoding in DA→NAc, we found that licking skew was highly correlated with DA→NAc skew (Fig. 5C; *r* = .77, *P* = .0002; Supplementary Table 1) and DA→NAc during ambiguous cues (Fig. 5D; *r* = .70, *P* = .001), but not DA→NAc during the positive or negative tones (Fig.5E,F; positive tone: *r* = .07, *P* = .8; negative tone: *r* = .07, *P* = .8). Similar to licking skew, DA→NAc skew was also stable across testing days (*r* = .72, *P* = .0008; Supplementary Table 1). Individual differences in global anticipatory licking rates (*r* = .08, *P* = .7; Supplementary Table 1) and licking rates during odor alone (*r* = .07, *P* = .77; Supplementary Table 1) did not predict DA→NAc skew, nor DA→NAc during ambiguous cues (*r* = .-.05, *P* = .9 and. *r* = .-.10, *P* = .7; Supplementary Table 1) These data indicate that trait-like affective bias is specifically encoded in DA→NAc responses to ambiguous cues, and that DA→NAc during ambiguous cues is not affected by individual differences in motivation and/or motor activity under these conditions.

**Figure 5.**
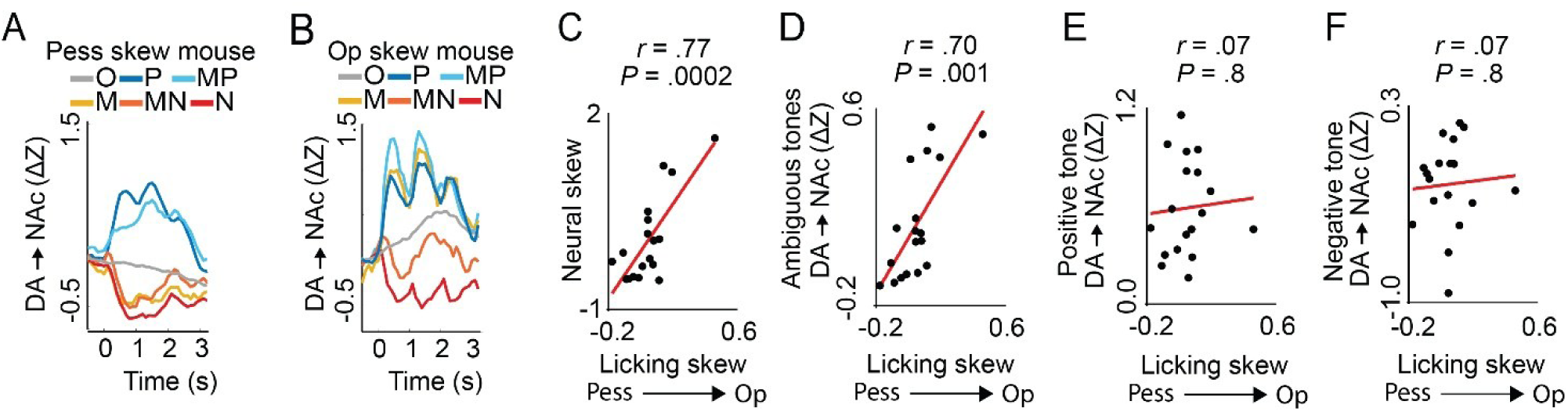
DA→NAc encodes trait-like pessimism during ambiguous cues. **(A)** Mean DA→NAc starting from tone onset in a mouse with a pessimistic licking skew. **(B)** Same as A but for a mouse with an optimistic licking skew. **(C)** DA→NAc skew vs licking skew (N = 18 learner mice). **(D)** Mean DA→NAc during ambiguous tones vs licking skew (N = 18 learner mice). **(E)** Mean DA→NAc during the positive tone vs licking skew (N = 18 learner mice). **(F)** Mean DA→NAc during the negative tone vs licking skew (N = 18 learner mice). C-F, each data point is from one mouse. Please see Supplementary Table 1 for more statistical information.

To more directly test if state and trait-like pessimism are encoded similarly in DA→NAc during ambiguous cues, we compared DA→NAc in mice with optimistic and pessimistic licking skews. We found that mice with pessimistic licking skews (licking skew < 0) had decreased DA→NAc during the ambiguous cues (Supplementary Figure 5A,B; *P* = .0012; Supplementary Table 1) but similar DA→NAc as mice with optimistic licking skews during the positive tone (Supplementary Figure 5A,B; *P* = .21; Supplementary Table 1) and negative tone (Supplementary Figure 5A,B; *P* = .996; Supplementary Table 1). Together, these results indicate similar encoding of state and trait-like pessimism in DA→NAc during ambiguous cues.

### Trait-like pessimism is associated with negatively shifted dopaminergic outcome signals

To determine if trait-like pessimism affected DA→NAc responses to reward and reward omission in a similar way as state pessimism, we compared DA→NAc in mice with optimistic and pessimistic licking skews during sucrose delivery and sucrose omission. Importantly, opposite of state pessimism, mice with pessimistic licking skews had a negatively shifted balance of positive and negative outcome encoding in DA→NAc (Fig. 6A, *P* = .01; Supplementary Table 1). This was due to significantly larger dips in DA→NAc during sucrose omission (Fig. 6B,D; LME, op vs pess mice, *P* = .01; LME, licking skew, *P* = .003; Supplementary Table 1), opposite of state pessimism, rather than a decrease in sucrose responses in DA→NAc (Fig. 6C,E; LME, op vs pess mice, *P* = 0.17; LME, licking skew, *P* = .18; Supplementary Table 1). The relationship between licking skew and DA→NAc during sucrose omission was not due to differences in neural activity just before the omission period, because the relationship remained when taking pre-omission activity into account with the LME (*P* = .005; Supplementary Table 1). These data indicate that state and trait-like pessimism affect DA→NAc similarly during ambiguous cues, but differently during outcomes, especially reward omission.

**Figure 6.**
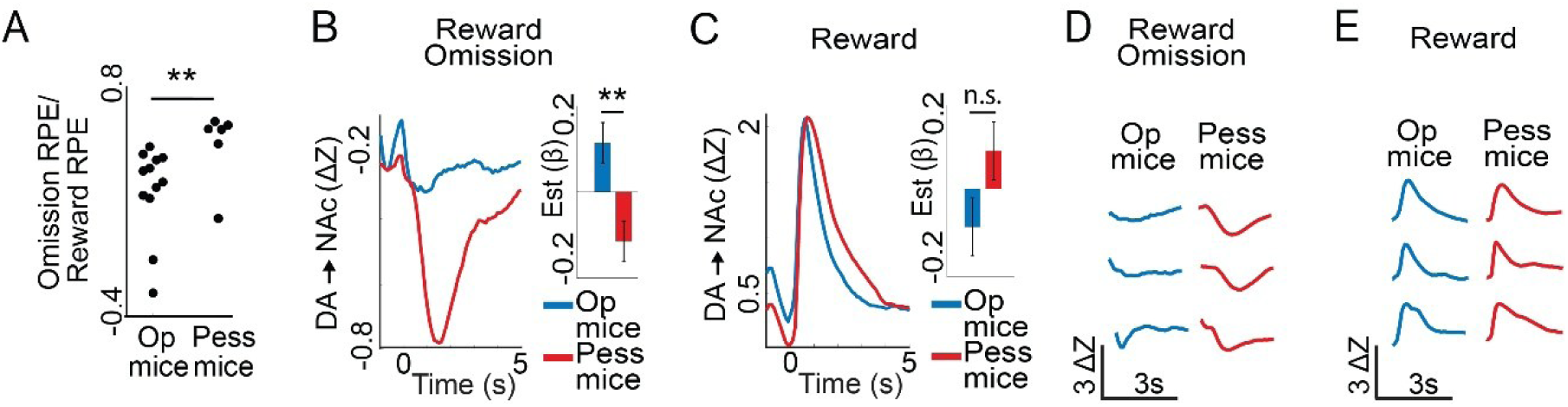
Negatively shifted balance of dopaminergic outcome signals in trait-like pessimism. **(A)** DA→NAc during sucrose omission relative to DA→NAc during sucrose consumption in mice with optimistic (N = 12) and pessimistic (N = 6) licking skews. The sign of the omission activity was inverted to make the ratios more intuitive. **(B)** Left, mean DA→NAc during sucrose omission (omission at 0 s) after tones for mice with pessimistic licking skews (“Pess mice”; N = 6) and optimistic licking skews (“Op mice”; N = 12). Right, corresponding beta estimates from LME which controls for possible correlations within mice and/or recording sites. **(C)** Same as B but during sucrose consumption. **(D)** Example mean DA→NAc responses to sucrose omission in 6 mice, as indicated. **(E)** Same as D but for sucrose consumption. ** P = .01. Please see Supplementary Table 1 for more statistical information.

### Trait-like pessimism is better explained by asymmetric RPE signaling than asymmetric plasticity

The DA→NAc outcome responses suggested that trait-like pessimism may be caused by asymmetric weighting of negative and positive dopaminergic RPE signals. To test this possibility, we simulated an asymmetric RPE signaling model and compared its predictions with those of an asymmetric plasticity model, another proposed mechanism of affective bias^28^ (Fig. 7A,B). In the asymmetric RPE signaling model, positive and negative outcome RPEs are differentially weighted in the DA signal itself^29^. In the asymmetric plasticity model, by contrast, DA RPE signals are weighted symmetrically, but positive and negative RPEs drive differentially weighted value updates downstream of DA signaling. Model simulations predicted identical positive relationships between positive-negative learning skew and ambiguous cue value (Fig. 7C), but different relationships between skew and the balance of positive and negative DA→NAc outcome RPEs (Fig. 7D). Thus, both models produce lower ambiguous cue values when skew is negative.

**Figure 7.**
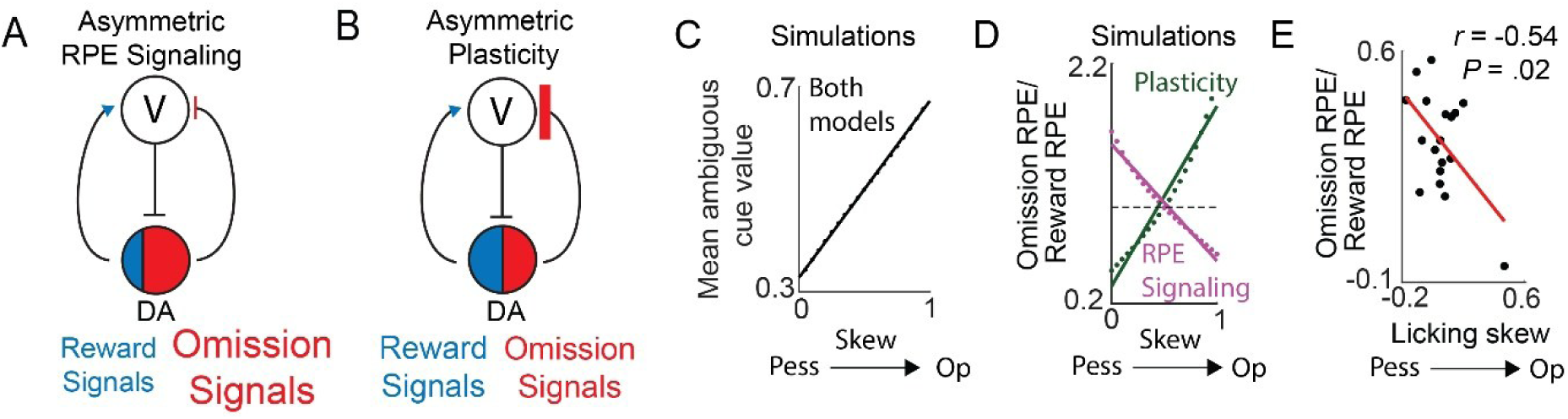
Asymmetric RPE signaling explains trait-like pessimism better than asymmetric plasticity. **(A)** Schematic of the asymmetric RPE signaling model of trait-like pessimism: heavily weighted omission RPEs relative to reward RPEs causes reduction in cue value (V). **(B)** Schematic of the asymmetric plasticity model of trait-like pessimism: equal weighting of omission and reward RPEs but more plasticity during omission RPEs than during reward RPEs causes reduction in cue value (V). **(C)** The asymmetric RPE signaling and asymmetric plasticity models predict identical mean ambiguous cue values relative to pessimistic/optimistic skew. **(D)** The asymmetric RPE signaling and asymmetric plasticity models predict different relationships between the balance of omission and reward RPEs and pessimistic/optimistic skew. Gray dashed line indicates expectation from standard TD model. **(E)** Negative relationship between the balance of omission and reward RPEs in DA→NAc and licking skew supports the asymmetric RPE signaling model (N = 18 learner mice).

However, the models make different predictions for outcome signaling in this scenario: the asymmetric RPE signaling model predicts relatively larger omission RPEs, whereas the asymmetric plasticity model predicts relatively smaller omission RPEs, because lower cue value would produce compensatory more positive outcome signaling, as observed during state pessimism. The empirical data matched the asymmetric RPE signaling model: negative licking skews among mice were associated with a more negative balance of DA→NAc reward and omission responses (Fig. 7E). Together, these data indicate that trait-like pessimism is maintained by a negatively shifted balance of positive and negative outcome signals due to greater weighting of reward omission RPEs.

We observed additional differences in mice with pessimistic licking skews. These mice showed more anticipatory licking during the negative tone (Supplementary Figure 5C,D; P = .03; Supplementary Table 1), resulting in weaker behavioral discrimination between the odor-only cue and the negative tone (Supplementary Figure 5G,H; P = .0008; correlation with licking skew, r = .63, P = .005; Supplementary Table 1), but not between the odor-only cue and the positive tone (Supplementary Figure 5I,J; P = 0.62; correlation with licking skew, r = .06, P = .82; Supplementary Table 1). Mice with pessimistic licking skews also had larger neural responses at odor onset (0–3 s; Supplementary Figure 5E,F; P = .03; correlation with licking skew, r = - 0.49, P = .04; Supplementary Table 1). However, the relationship between licking skew and reward omission RPEs remained significant when anticipatory licking, odor-onset neural responses, or both were included as co-regressors in the statistical model (LME, with odor-onset neural response included, P = .0007; with anticipatory licking included, P = .005; with both included, P = .005; Supplementary Table 1). Thus, enhanced reward omission RPEs were not explained by these additional behavioral or neural correlates of trait-like pessimism. Interestingly, there were no differences in DA→NAc neural discrimination between mice with pessimistic and optimistic skews, neither for the odor-only/positive tone discrimination (Supplementary Figure 5M,N; *P* = 0.49; correlation with licking skew, *r* = .12, *P* = .65; Supplementary Table 1) nor odor-only/negative tone discrimination (Supplementary Figure 5K,L; *P* = 1; correlation with licking skew, *r* = -.07, *P* = .77; Supplementary Table 1), despite the difference in behavioral discrimination between odor-only/negative tone trials (Supplementary Figure 5G,H).

The discrepancy between odor-only/negative tone behavioral discrimination (worse than optimists) and odor-only/negative tone DA→NAc discrimination (same as optimists) in mice with pessimistic licking skews indicated that they discriminated the odor-only and negative cues similarly to mice with optimistic licking skews but lacked behavioral inhibition or response flexibility during the negative tone. Consistent with behavioral inflexibility in mice with pessimistic licking skews, they had shorter duration licking bouts during sucrose consumption that did not decrease within sessions like those of mice with optimistic licking skews (Supplementary Figure 5O; LME, op vs pess mouse x Trial Number, *P* = 2.7 × 10^−5^; Supplementary Table 1). Mice with pessimistic licking skews also did not decrease licking during the positive tone within sessions as much as mice with optimistic licking skews (Supplementary Figure 5P; op vs pess mouse x Trial Number, *P* = 1.4 × 10^−5^; Supplementary Table 1). Thus, these data suggest a link between behavioral inflexibility and pessimistic cue interpretation, traits that have previously been independently linked to stress-related psychiatric disorders^26,36,37^.

## Discussion

Here we use a Pavlovian ambiguous-cue task in mice to show that trial-to-trial, state pessimism is encoded via reduced DA→NAc during ambiguous cues and a compensatory increase in relative DA→NAc responsiveness to reward compared to reward omission. We also find that individual differences in trait-like pessimistic anticipatory licking are encoded similarly as state pessimism during ambiguous cues but have an opposite balance of reward and reward omission signaling. Thus, these two forms of pessimism have partially dissociable dopamine dynamics.

Although anticipatory licking can be affected by attention, motivation, and trial engagement, the relationships between DA→NAc and pessimism could not easily be explained by these factors. State pessimism was associated with reduced DA→NAc during the ambiguous cues, not just lack of DA→NAc response, and only in mice that learned the motivational significance of the cues. Trait-like pessimism was associated with decreased DA→NAc during ambiguous cues and neither licking bias nor DA→NAc was related to individual differences in global anticipatory licking. Furthermore, trait-like bias did not change systematically within sessions over time, unlike anticipatory licking, and was stable across testing sessions. Instead, the data indicate that DA→NAc during ambiguous cues and outcomes is strongly affected by subjective state and trait-like affective bias.

Several observations suggested behavioral inflexibility in pessimistic mice: selectively more licking during the negative tone despite similar DA→NAc during the negative tone, and no decrease in licking during the positive tone nor lick bout duration during sucrose delivery within sessions. These mice also had larger phasic DA→NAc during odor onset. It is unclear how trait-like pessimism relates to these observations mechanistically; however, it is notable that anxiety has been linked to compulsive behaviors, sensory hypersensitivity, and negative feedback hypersensitivity^36,38–40^. Therefore future studies that examine the relationship between trait-like pessimism, DA→NAc, and measures of or pharmacological treatments for anxiety may lead to a mechanistic understanding of these links.

Our observations during state pessimism are consistent with TD reinforcement rules^1,41^ and compensatory changes in learning that depend on subjective reward expectation. The complementary pattern during state optimism illustrates a potential computational constraint on transiently shifting expectations in a positive direction. More optimistic interpretations of ambiguous cues were followed by a negatively shifted balance of outcome-related DA→NAc signals. Thus, changing an expectation without changing subsequent outcomes generates teaching signals that oppose the initial bias. This observation may provide a reinforcement-learning framework for understanding why unrealistically positive expectations are not invariably associated with greater long-term well-being^42–45^.

Unlike the compensatory outcome responses observed during transient state bias, trait-like pessimism was associated with the opposite balance of outcome-related DA→NAc activity, due to larger omission RPEs. This dissociation suggests that state and trait-like pessimism are not simply the same process measured over different timescales but instead arise from distinct neural mechanisms.

Our findings favor asymmetric RPE signaling as a mechanism for trait-like pessimism, rather than asymmetric plasticity downstream of dopamine signaling^28^, and are consistent with recordings demonstrating asymmetric RPE signaling in individual midbrain DA neurons^29^. However, it remains to be determined how a population-level asymmetry in individual mice arises—for example, via a population-wide shift toward more negative RPE asymmetry or increased abundance of neurons with negative asymmetry—and whether it is generated by synaptic or cellular mechanisms within the VTA^2,46–51^ or upstream regions that are implicated in mood disorders, such as the lateral habenula^52–66^.

## Methods

### Animals

A total of 28 adult 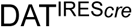 heterozygotes (DAT-Cre or B6.SJL-Slc6a3tm1.1(cre)Bkmn/J mice; The Jackson Laboratory; 006660;17 male, 11 female) were used in this study. Animals were singly housed on a 12 hr dark/12 hr light cycle (light from 06:00 a.m. to 06:00 p.m.). All procedures were approved by the Institute Animal Care and Use Committees of the University of Texas Southwestern Medical Center.

### Surgery

Mice (2-4 months old) were anesthetized with isoflurane (1-2% at 0.5 - 1.0 L/min) and given analgesia (carprofen, 5 mg/kg, I.P) before the surgery, and for 3 days following the surgery. Mice were also given antibiotics dissolved in water (sulfadiazine/trimethoprim oral suspension, or sulfatrim, 1 mL/400 mL vol/vol in water) post-surgery for 1 week.

Calcium indicators were injected into the ventral tegmental area (VTA, AP: −3.1 ML: ±1.35 DV:-4.16, −4.47), and optical fibers were implanted into the nucleus accumbens core (NAc, AP: +1 ML: ±1.84 DV: −4.26, with 5 degree angle medially). These injections and implants were bilateral in one group of 11 mice, and unilateral and counterbalanced in another group of 16 mice. Calcium indicators used were cre-dependent RCaMP1b (AAV2/1.Syn.Flex.NES-jRCaMP1b, Addgene #100850) in the first group, and GCaMP8f (AAV2/5-Syn-FLEX-jGCaMP8f, Addgene # 162379) in the other group. Viruses were injected with a volume and concentration of 250 nL and 5×10^12^ GC/mL per injection. Optical fibers used for implantation had a core diameter of 200 μm (RWD; R-FOC-BL200C-39NA). Custom-designed head-bars (Hayes Manufacturing) were attached to the skull using dental cement (C&B Metabond). All experiments were conducted at least 3 weeks after surgery to allow virus expression and recovery.

### Ambiguous-cue paradigm

Head-fixed mice were first trained to associate a sucrose reward (∼ 5 uL; 10% sucrose wt./vol in water) with an olfactory cue (vaporized 4% isoamyl acetate in mineral oil) presented for 6.2 seconds (Stage 1). Sucrose was delivered immediately after the odor offset through a spout placed in front of the mouse’s mouth, and licking was measured using an optical lickometer (Optical Lickometer, Product ID 1020, Sanworks).

We confirmed that mice learned the odor/reward association by measuring anticipatory licking during the odor. Mice with anticipatory licking in at least 85% trials for at least two consecutive sessions were moved to the next stage of training. These mice were then given 1-3 sessions of odor trials with 50% probability of reward (Stage 2).

Mice were then trained to associate a negative and positive tone (3 or 12 kHz, counterbalanced) with 0% and 100% reward, respectively (Stage 3). The tones were pipped and presented from 3 to 6.2 seconds from the onset of odor. The tones were on for 200 ms and off for 800 ms, for 4 cycles. Mice that showed a significant difference in licking for negative and positive tones (t-test, P < 0.01) were categorized as tone learners (“learners”; N = 19 mice). The rest were categorized as tone non-learners (“non-learners”; N = 9 mice). All mice received at least 15 sessions of tone training. One learner mouse was excluded because of a lack of DA→NAc signal.

All mice, regardless of whether or not they learned the associations to the training tones, were presented with ambiguous tones of frequencies intermediate to the positive and negative tones (5.25, 7.5 and 9.75 kHz). The probabilities of occurrence of each of the ambiguous tones, training tones, and odor trials were 4%, 22% and 44%, respectively. The ambiguous cues were followed with a probability of reward scaled to their acoustic proximity to the negative and positive tones (mid negative 25%, mid 50% and mid positive 75%). This was to ensure that the mice did not stop responding to the ambiguous cues over time.

All trials during training and testing sessions were randomized using a pseudo-random number generator (Arduino) and a new random seed for each session. Inter-trial interval lengths (10-30 seconds) were also randomized.

### Fiber photometry

Fiber photometry was performed according to the Neurophotometrics manual and published protocols^67^. A fiber-optic patch cord (Doric Lenses) was attached to the implanted optical fiber with cubic zirconia sleeves covered with black, shrinkable tubing (Sleeve_ZR_1.25-BK, Doric Lenses). The other end of the fiber-optic cable was coupled to a Neurophotometrics LED port. Bonsai was used to control the system and 560 or 470 and 415 nm LEDs. Light at the fiber tip ranged from 50 to 100 μW. LED frequencies were either 40 Hz, with 10Hz for each of the LEDs and for LED off state, or down-sampled/interpolated to 40 Hz to maintain uniformity across all mice and sessions.

### Data analysis

Fiber photometry: Calcium indicator signal was smoothed using a span of 20,000 data-points to account for photobleaching and subtracted from the original, non-smoothed data, then Z-transformed using the mean of each session, and baseline subtracted (time period of 10s preceding onset of odor cue). Z-scored neural data were aligned with behavioral data using TTLs sent from Arduino to Cheetah, using Arduino scripts.

Optimistic/pessimistic trial analysis using surrounding odor trials: Tone trials were separated into pessimistic and optimistic categories by computing the average of licking during the last second of odor presentation (5.3 to 6.2s from odor onset) in two odor-only trials before the trials of interest and two odor-only after them (Av_odor_). The licking during tone presentation on the current tone trial for the same time interval was also computed (Current). If Current < Av_odor_, that tone trial was categorized as ‘pessimistic’, else if Current > Av_odor_ the trial was categorized as ‘optimistic’ (Figure 2A). Statistical analyses of differences between pessimistic and optimistic trials during tone presentation were done on data averaged from 3 to 6.2s from odor onset for each trial (i.e., when the tones were presented). Statistical analyses of differences between pessimistic and optimistic trials during outcomes were done on average neural data within a 5 second window from sucrose delivery or omission. Additionally, for sucrose consumption analyses, only trials in which the mouse licked within 0.5 seconds of sucrose delivery were included, to ensure alignment of neural activity to sucrose consumption.

Paired mouse analysis: Mice with the same trial histories during training and testing sessions were paired during analysis. Trials that co-occurred with pessimistic and optimistic reference trials (‘Pess Ref’ and ‘Op Ref’, respectively) in the paired mouse were then categorized as paired pessimistic and optimistic trials (‘Pess Paired’ and ‘Op Paired’). Data from 7 pairs (14 mice, including 2 pairs of learners, 2 pairs of non-learners and 3 mixed pairs) were used.

Skew calculations: Skews were calculated for licking and neural activity for each mouse by calculating the difference between actual mean licking/neural activity in ambiguous-tone trials (during the tone periods - 3 to 6.2 seconds from odor onset) and values obtained by linear interpolation from mean licking/neural activity in negative and positive tone trials (see Figure 4A). Licking skews > 0 were called optimistic, and licking skews < 0 were called pessimistic. Licking skew values were divided by 10 to effectively keep them between −1 and 1. Statistical analyses of differences in licking and neural activity in optimistically and pessimistically skewed mice during tone and outcome presentation were done using the same time periods used for optimistic/pessimistic trial analysis.

AUROC for odor-only vs positive/negative tone licking and neural discrimination: AUROC was calculated as the probability that each positive/negative tone value (licking or neural during the tone interval) was greater than each odor-only value (licking or neural during the same time interval). For the negative tone, to make the graph more intuitive, this probability value was inverted so that a value of 1 means all negative tone values were less than all odor-only values.

Lick bout duration during sucrose consumption: Licking during the sucrose consumption interval (0-5 seconds from sucrose delivery) was separated into bouts if there was a minimum of 0.1 seconds pause in licking. The duration of these licking bouts was then averaged for each trial and only bouts > 100 ms were included in the average.

### Model simulations

We used simple RPE-like computational models to determine whether different mechanisms of positive-negative outcome asymmetry could account for the relationship between trait-like licking skew, ambiguous cue value, and DA→NAc responses during reward and reward omission. Cues were assigned reward probabilities matching the task structure: near-negative, 25%; midpoint, 50%; and near-positive, 75%. For each cue, the raw outcome prediction error was defined as δ = r − V, where r was the outcome value, set to 1 for sucrose delivery and 0 for sucrose omission, and V was the cue value. To incorporate trial-to-trial variability in value representation, cue values were sampled from Gaussian distributions centered on their stable values.

Stable cue values were computed by solving for the value at which the expected learning update was zero. Positive updates were multiplied by (Vmax − V)^0.5^, whereas negative updates were multiplied by (V − Vmin)^0.5^, where Vmin and Vmax were the lower (0) and upper (1) value bounds. Thus, positive updates diminished as cue value approached the upper bound, and negative updates diminished as cue value approached the lower bound for all models.

We compared three models. In the standard TD model, positive and negative prediction errors were represented and learned from symmetrically. In the asymmetric plasticity model, the DA-like RPE signal was not itself weighted asymmetrically, but positive and negative RPEs drove value updates with different gains. Thus, for δ ≥ 0, the learning update was α^+^ _*_ δ, whereas for δ < 0, the learning update was α^−^ _*_ δ. In the asymmetric RPE signaling model, positive and negative RPEs were differentially weighted in the expressed DA-like signal, and the weighted signal was also used for learning. Thus, for δ ≥ 0, the signal/update was k^+^ _*_ δ, whereas for δ < 0, the signal/update was k^−^ * δ. For both asymmetric models, the relative strength of positive versus negative weighting was varied continuously using a skew parameter ranging from 0 to 1, where 0 meant α^−^ or k^−^ was three times larger than α^+^ or k^+^ and 1 meant α^+^ or k^+^ was three times larger than α^−^ or k^−^. Therefore, low skew values corresponded to stronger negative than positive weighting, whereas high skew values corresponded to stronger positive than negative weighting.

For each model and skew value, we computed stable values for each cue and then sampled and averaged the outcome-period RPEs. Outcome-period RPEs were defined as sucrose delivery (1) minus cue value for reward responses and omission (0) minus cue value for omission responses, and these raw RPEs were weighted by k^+^ or k^−^ in the asymmetric RPE signaling model, then averaged across the near-negative, midpoint, and near-positive cues. The balance of positive and negative outcome RPE signaling was quantified as the absolute value of the mean ambiguous omission RPE divided by the mean ambiguous reward RPE.

### Histology

To confirm the position of recording sites, and to perform immunohistochemistry, mice were given an anesthetic overdose (ketamine-dexmedetomidine, 0.2 mL) and perfused with phosphate buffered saline (PBS), followed by paraformaldehyde (4% wt./vol in PBS). Brains were post-fixed for 24 h. Whole brains were then transferred to 30% sucrose with 0.02% sodium azide, until they were sectioned (40 um thickness) using a cryostat (CM 3050 S, Leica Biosystems). The fiber tracks were mapped onto standard atlas sections by visual inspection (Paxinos and Franklin, 2001). Recording sites are shown in Figure S1A.

### Immunohistochemistry

Slices were washed in PBS, then immunolabeled with a primary Anti-Tyrosine Hydroxylase Antibody (TH, EMD Millipore Sigma MAB318; 1:2000; Lot 3083054; with 0.2% Triton and 3% normal goat serum) for 3 days at 10 °C, then secondary immunolabeling for 3 hours at room temperature (1:300 goat-anti-mouse 647), followed by washing and mounting with Vectashield Anti-fade with DAPI). High-resolution (1024×1024), tiled images were taken with a 63X objective and Zeiss LSM 800 confocal microscope (14 pictures of 4 slices from 2 mice).

### Statistics

Except where indicated otherwise, we used linear mixed-effects models (LME)^34^ in MATLAB using the fitlme function. This method accounts for possible correlations among data coming from the same optical fiber and animal by modeling them as random effects. This method is usually more conservative than other statistical tests that assume statistical independence for data from the same recording site and/or animal. As indicated in the text and Supplementary Table 1, in some cases, other fixed effect regressors were used (such as the trial (tone) type to factor out the objective trial type when testing for subjective bias). Post-hoc linear hypothesis tests were done to probe specific comparisons based on ANOVA results from the LME. All other statistical analyses were done in MATLAB or GraphPad Prism.

## Acknowledgments

We thank Daisuke Hattori, Todd Roberts, and Peter Tsai for helpful discussion. Funding for this project was provided by UT Southwestern Medical Center and the NIH.

