## Supplementary Figures for "Dopaminergic signatures of state and trait-like pessimism in mice"

Supplementary Figure 1

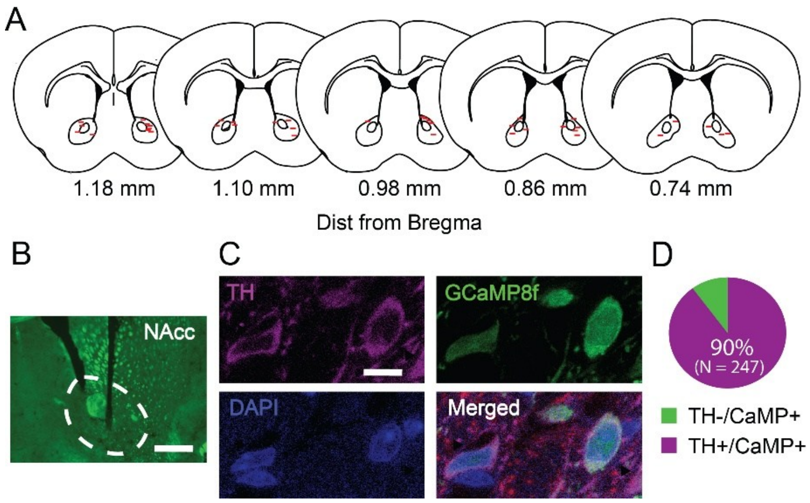


**Supplementary Figure 1. Placement of optical fibers and co-localization of calcium indicators and tyrosine hydroxylase. (A)** Location of optical fibers mapped onto standard atlas sections (Paxinos, G and Franklin, K.B.J. The mouse brain in stereotaxic coordinates, 2001). N = 38 sites from 27 mice. **(B)** Example of a fiber tract in the NAc core (NAc). Scale bar, 750 µm. **(C)** Immunohistochemistry images showing the localization of tyrosine hydroxylase (TH), GCaMP and DAPI in two example neurons. Scale bar, 10 µm. **(D)** Quantification of calcium indicator-expressing neurons expressing TH. Data from 247 cells in 2 mice.

Supplementary Figure 2

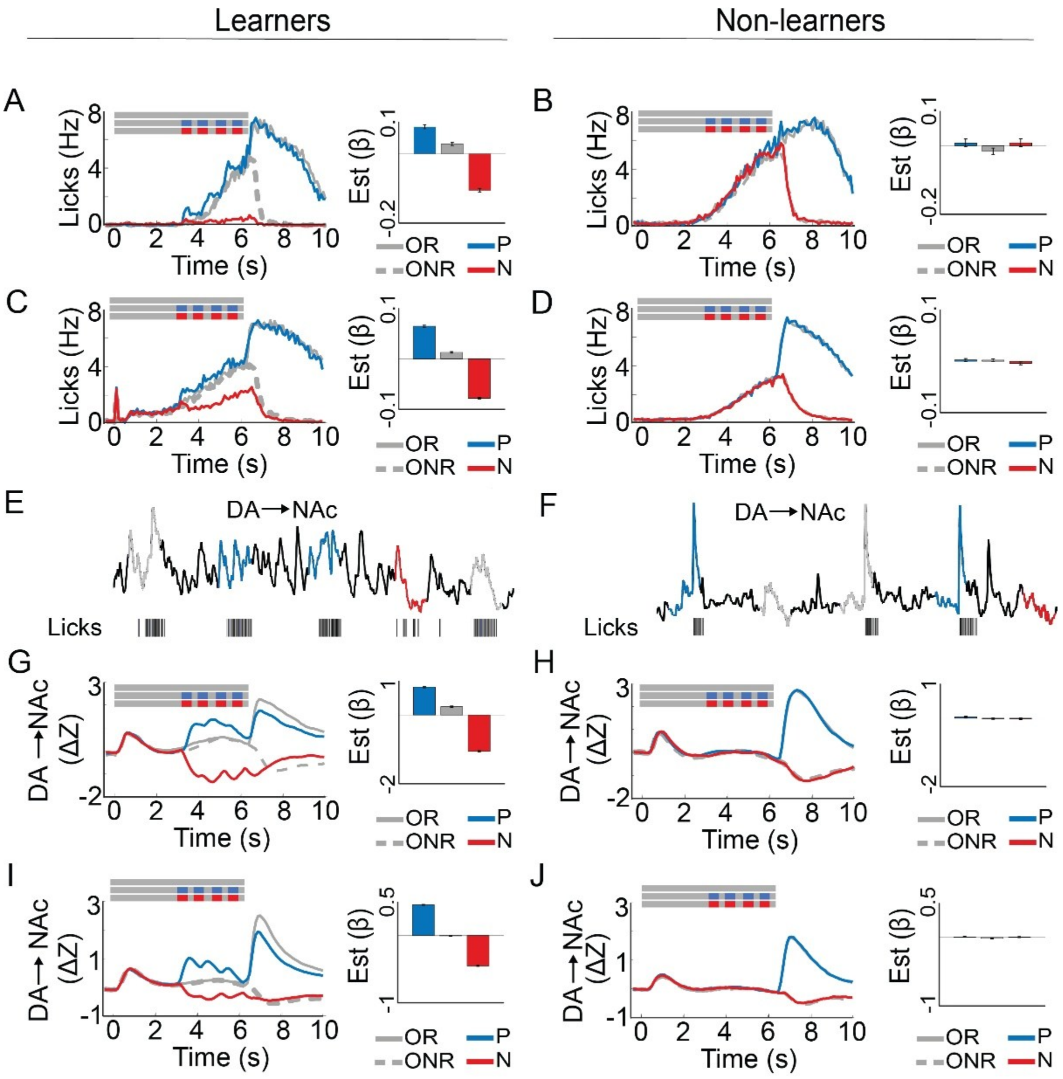


**Supplementary Figure 2. Licking and DA→NAc in learners and non-learners.** (**A-J**) Left, mean licking or DA→NAc activity. Right, corresponding beta estimates and standard error from linear mixed-effects modeling (LME, see Methods). The column labels, learners (18 mice) and non-learners (9 mice), refer to all data in A-J. **(A)** Licking after training in an example learner mouse. The three gray bars on the top left denote odor, positive and negative tone onset in the respective trial types. OR, odor-reward trials. ONR, odor-no reward trials. P, positive tone trials. N, negative tone trials. **(B)** Same as A, but for a non-learner. **(C)** Mean licking after training in learners. Abbreviations as in A. **(D)** Same as C, but for non-learners. **(E)** Raw DA→NAc in five consecutive trials in an example learner. Ticks below indicate individual licks. Colors as in A-D. **(F)** Same as E, but in a non-learner. **(G)** DA→NAc after training in an example learner. Abbreviations as in A. **(H)** Same as G, but in a non-learner. **(I)** Mean DA→NAc after training in learners. Abbreviations as in A. **(J)** Same as I, but in non-learners.

Supplementary Figure 3

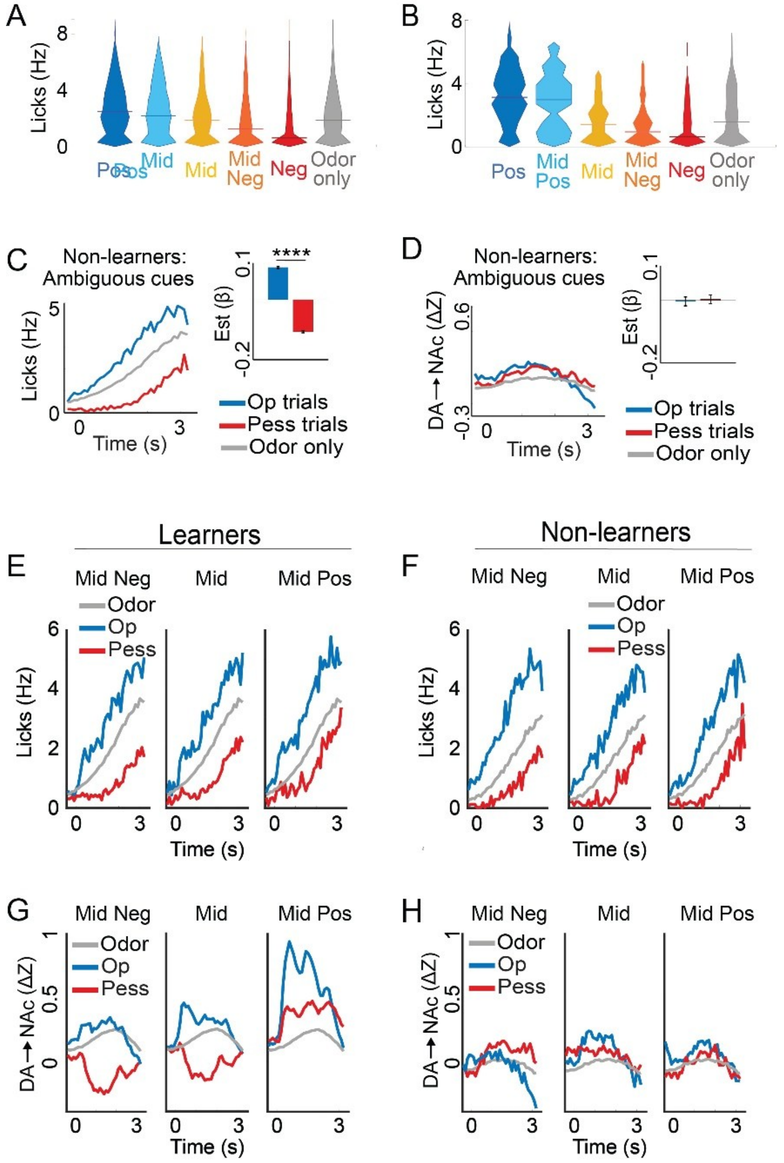


**Supplementary Figure 3. Licking and DA→NAc during state affective bias in learners and non-learners. (A)** Violin plots indicating the distribution of licking during individual trials in learners. Lines indicate medians. **(B)** Same as A, but in an individual mouse. **(C)** Left, mean anticipatory licking during odor-only trials (Odor) and optimistic (Op) and pessimistic (Pess) ambiguous cue trials in non-learners. Right, corresponding beta estimates from LME which controls for objective trial type and possible correlations within mice and/or recording sites. **(D)** Same as C, but DA→NAc instead of licking. **(E)** Same as C, but for individual ambiguous tone types in learners. **(F)** Same as E, but in non-learners. **(G)** Same as E, but DA→NAc instead of licking. **(H)** Same as G, but in non-learners. ****P < 0.0001. Please see Supplementary Table 1 for more statistical information.

Supplementary Figure 4

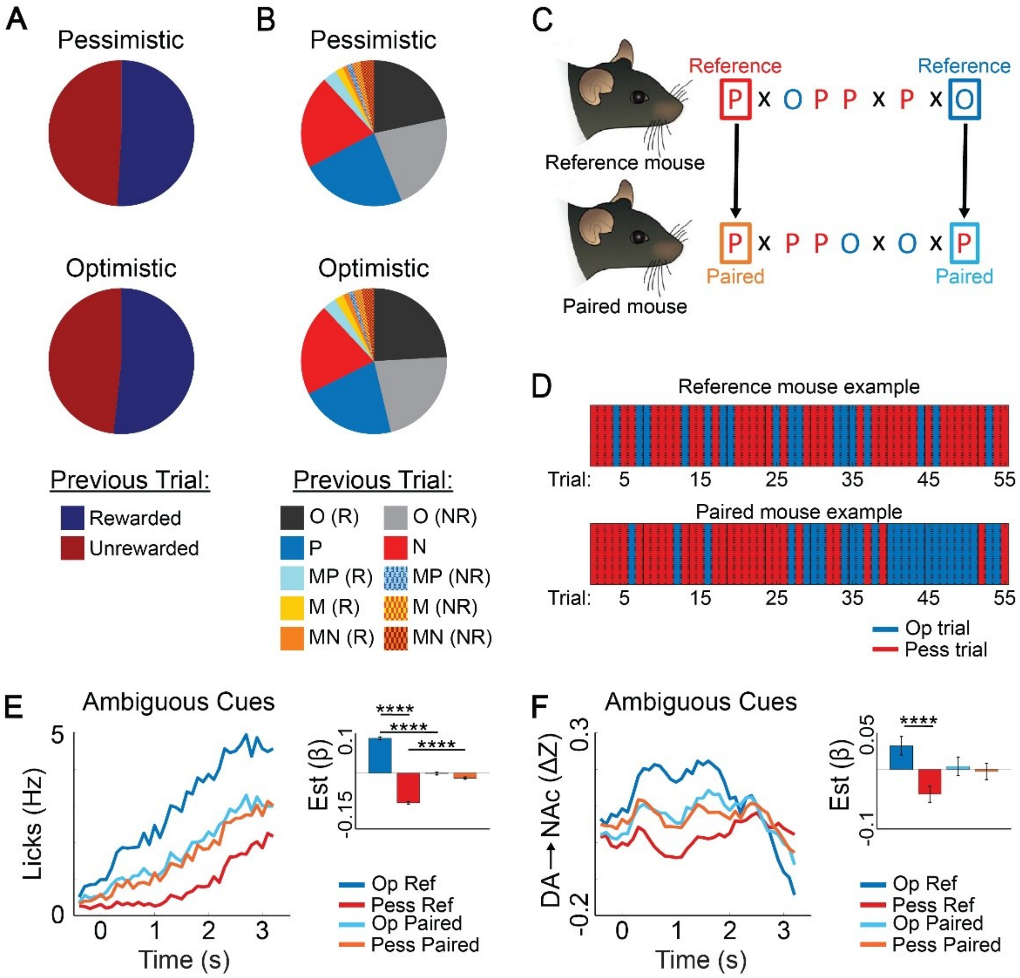


**Supplementary Figure 4. State pessimism is subjective. (A)** Pie chart showing the proportion of immediately preceding trials that are rewarded or unrewarded for pessimistic and optimistic trials. (N = 9885 trials from 18 mice; chi-square, n.s.). **(B)** Pie chart showing the proportion of immediately preceding trials that are of each type for pessimistic and optimistic trials. N same as in A. chi-square test, n.s. Trial-type abbreviations as in Figure 1. R, rewarded. NR, not rewarded. **(C)** Schematic of the method used to compare trials in mice with identical training and testing histories. P, pessimistic trials. O, optimistic trials. X, odor-only trials. **(D)** Examples of optimistic/pessimistic mid-tone trial classification for a pair of learner mice with identical trial orders. Trials shown in chronological order. **(E)** Left, Anticipatory licking during ambiguous cues. Right, beta estimates from LME in reference (Ref) mice and their corresponding paired (Paired) mice that accounts for objective trial type. Optimistic (Op) and pessimistic (Pess) refers to classification of Ref mouse behavior only (N = 7 mouse pairs). **(F)** Same as E, but DA→NAc instead of licking. ****P < 0.0001. Please see Supplementary Table 1 for more statistical information.

### Supplementary Figure 5

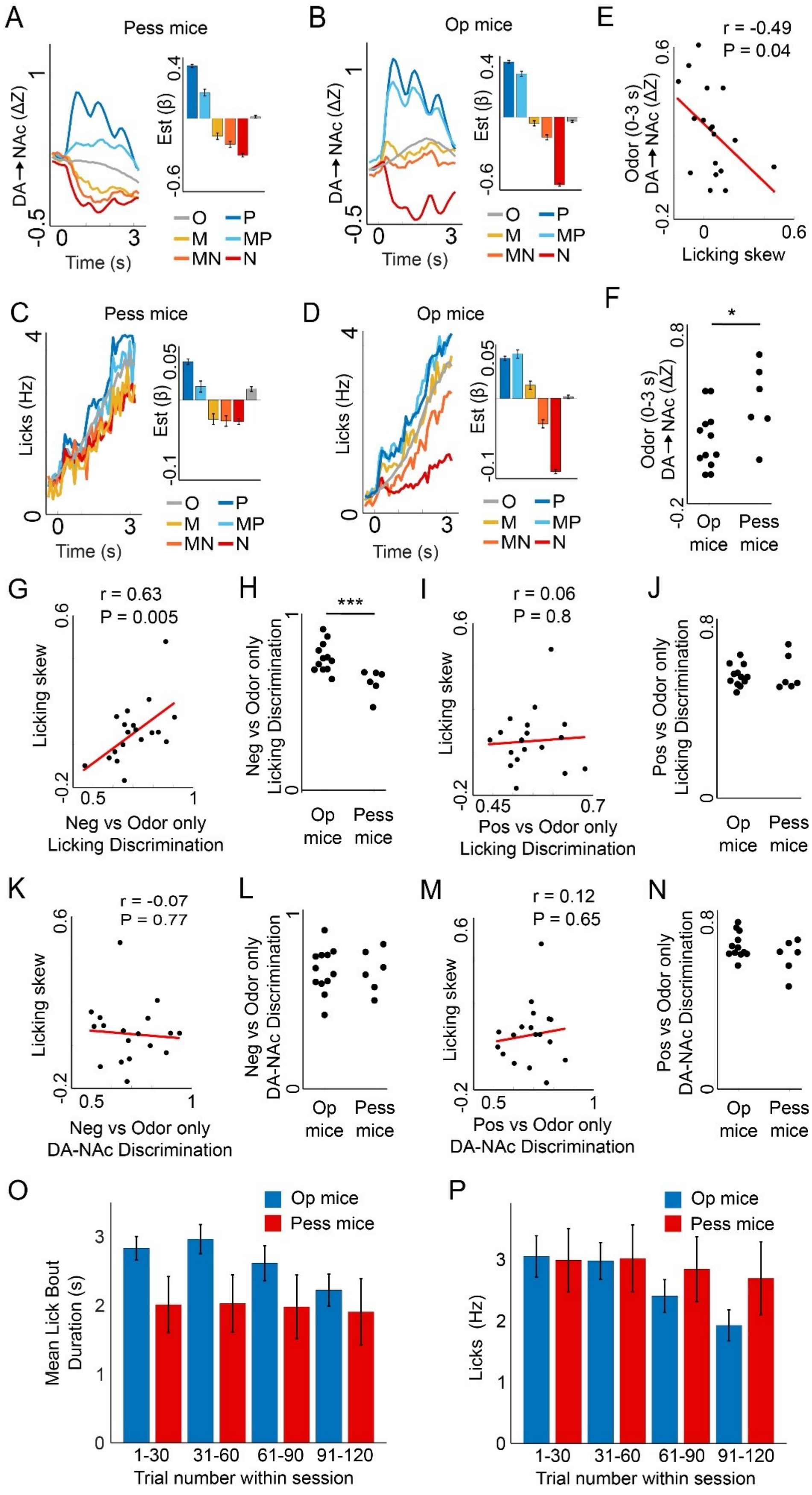

**Supplementary Figure 5. Licking and DA→NAc in mice with optimistic and pessimistic licking skews. (A-D)** Left, mean DA→NAc or anticipatory licking during tones in the test sessions. Right, corresponding beta estimates from LME which controls for possible correlations within mice and/or recording sites. O, odor only. P, positive tone. MP, mid-positive tone. M, mid-tone. MN, mid-negative tone. N, negative tone. **(A)** DA→NAc in mice with pessimistic licking skews (N = 6 mice). **(B)** DA→NAc in mice with optimistic licking skews (N = 12 mice). **(C)** Licking in mice with pessimistic licking skews (N = 6 mice). **(D)** Licking in mice with optimistic licking skews. (N = 12 mice). **(E)** Licking skew vs DA→NAc during the first 3 seconds of the odor cue (N = 18 mice). **(F)** DA→NAc during the first 3 seconds of the odor cue in mice with optimistic and pessimistic licking skews (N = 12 and 6 mice). **(G)** AUROC for discriminative licking between the negative tone and odor-only trials vs licking skew (N = 18 mice). **(H)** AUROC for discriminative licking between the negative tone and odor-only trials for mice with optimistic and pessimistic licking skews (N = 12 and 6 mice). **(I)** AUROC for discriminative licking between the positive tone and odor-only trials vs licking skew (N = 18 mice). **(J)** AUROC for discriminative licking between the positive tone and odor-only trials for mice with optimistic and pessimistic licking skews (N = 12 and 6 mice). **(K)** AUROC for discriminative DA→NAc between the negative tone and odor-only trials vs licking skew (N = 18 mice). **(L)** AUROC for discriminative DA→NAc between the negative tone and odor-only trials for mice with optimistic and pessimistic licking skews (N = 12 and 6 mice). **(M)** AUROC for discriminative DA→NAc between the positive tone and odor-only trials vs licking skew (N = 18 mice). **(N)** AUROC for discriminative DA→NAc between the positive tone and odor-only trials for mice with optimistic and pessimistic licking skews (N = 12 and 6 mice). **(O)** Mean lick bout duration during sucrose consumption within sessions in mice with optimistic (Op) and pessimistic (Pess) licking skews. LME, op vs pess mouse x Trial Number, $P = 2.7 \times 10^{-5}$. **(P)** Mean lick rate during positive tone trials within sessions in mice with optimistic (Op) and pessimistic (Pess) licking skews. op vs pess mouse x Trial Number, $P = 1.4 \times 10^{-5}$. Pearson's correlation used in E,G,I,K,M. Mann-Whitney test used in F,H,J,L,N. $^{*}P < 0.05$, $^{***}P < 0.001$. Please see Supplementary Table 1 for more statistical information.
